# Genotyping of *Orientia tsutsugamushi* circulating in and around Vellore (South India) using TSA 56 gene

**DOI:** 10.1101/2022.12.29.522276

**Authors:** Janaki Kumaraswamy, Punitha Govindasamy, Lakshmi Surya Nagarajan, Karthik Gunasekaran, KPP Abhilash, John Antony Jude Prakash

**Author notes:** **Corresponding author** Dr. John Antony Jude Prakash, Professor and Head, Department of Clinical Microbiology, Christian Medical College, Vellore, Tamil Nadu, India.

## Abstract

The immunodominant TSA 56 gene of *Orientia tsutsugamushi*, (scrub typhus agent) has four variable regions (VD-I to VD-IV) making it useful for genotyping. As of date the genotyping data from India is based on partial 56kDa gene sequence analysis. The complete TSA 56 gene sequence is important for knowing the circulating strains and for designing region specific diagnostics and vaccines. This study was undertaken to determine *Orientia tsutsugamu*s*hi* genotypes circulating in and around Vellore using complete and partial TSA 56 gene. Of the 379 whole blood samples from suspected scrub typhus patients, 162 were positive by 47 kDa qPCR. Long protocol to amplify the complete TSA 56 gene (≈1605 bp) was performed on 21 samples. On the same 21 samples the partial gene sequence was also amplified using the Horinouchi (≈650bp) and the Furuya (≈480 bp) protocol. Using a combination of Sanger and Nanopore technology complete sequence was obtained for 9 and near complete (1551 to 1596 bp) for 4 respectively. As Furuya protocol gave multiple bands we obtained 480 bp sequences from the 13 complete gene sequences by *in silico* analysis. In contrast, 650bp sequences were obtained for 11 samples while for the remaining two we derived the 650 bp sequences from the complete gene sequences (Long protocol). Phylogenetic analysis of the complete gene (Long protocol) which includes VD-I to VD-IV region and partial gene (Horinouchi) which amplifies the VD-I to VD-III regions showed identical genotypes. Twelve belonged to TA763 genotype and one belongs to Karp genotype. The Furuya sequence (*in silico*) correctly identified the Karp genotype and 10 of the TA763 genotypes. Two TA763 genotypes (identified by complete and 650 bp partial gene analysis) were misidentified by Furuya sequence analysis as Karp genotype.

The limited analysis showed the commonest *Orientia tsutsugamushi* genotypes circulating in and around Vellore is TA763 and that the 650 bp (Sanger) sequencing could be a cost effective method for identifying the scrub typhus genotypes. However, these results need to be validated by larger prospective multi-centric studies.

## Introduction

Scrub typhus is a vector borne acute febrile illness caused by *Orientia tsutsugamushi* (formerly known as *Rickettsia tsutsugamushi*) and it is common in India (1). The etiological agent is transmitted to rodent or human by bite of the infected chiggers (larval) of trombiculid mite (2). Till recently, scrub typhus was thought to be endemic only in the tsutsugamushi triangle (3). There is evidence now of scrub typhus like infections from the Middle East (Dubai), Africa (South Africa), Europe (France) and Chile in South America (4).

*Orientia tsutsugamushi* has many antigenic variants and >40 antigenic strains (5) and virulence has been attributed to regional strain differences, suggesting that virulence of *Orientia tsutsugamushi* is related to genotype (6). Genotypic classification of *Orientia tsutsugamushi* is based on the variations in the immuno-dominant outer membrane protein, the 56-kDa type-specific antigen (7). Though all individuals with scrub typhus have antibodies to the 56 kDa antigen, immunity is strain (genotype) specific. The immunogenicity of this antigen has made it a good diagnostic and vaccine candidate (8).

The gene which encodes the 56 kDa protein (TSA 56 gene) consists of an open reading frame (ORF) of ≈1600 bp encoding 530 amino acids, and has four variable domains VD I to IV, which are described in Figure 1 (9,10). The variations in these four domains has led to the observation of various of genotypes like Karp, Kato, Gilliam, Boryong, Kawasaki, Shimokoshi, TA763 and TA686 (10,12,13). Data on the complete TSA 56 gene is available from Thailand, Laos, South Korea, Malaysia, Taiwan, China, Japan, Vietnam, Myanmar, Cambodia and New Guinea (10, 14-16).

**Figure 1:**
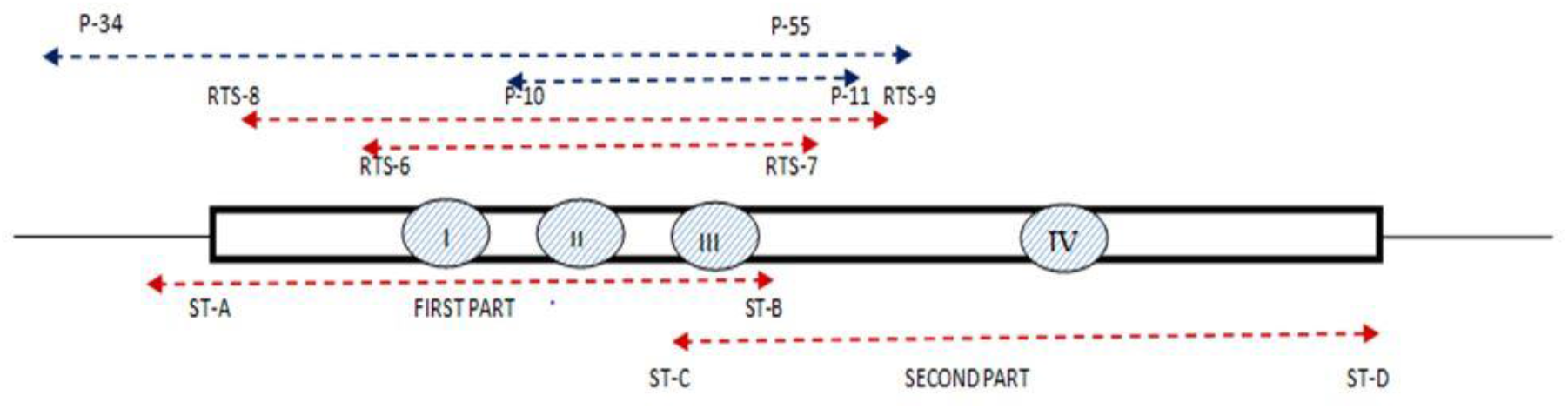
Binding site of primers used for 56kDa gene amplification.

From India, only few reports are available on *Orientia tsutsugamushi* genotypes but they are based on partial TSA 56 gene sequence which covers two of the four variable domains. Most of the studies are from North, South and North East India. The predominant genotypes reported Kato, Karp, Kato like, Karp like, and Gilliam strains (11). In Northern India, Boryong (63.4 %) is the predominant strain, followed by Karp (23.6%), Gilliam (11.8%) and Kawasaki (1.2 %) whereas in South and North-East India, Kato is the most predominant type followed by Karp like and Gilliam strain (11,12). A study on chiggers collected from Hosur and Vellore, identified TA716 and Kato genotype based on Horinouchi protocol (13). Chunchanur *et al*., (2019) found Gilliam strain in Karnataka, India (14). Studies from Andhra Pradesh, Madhya Pradesh and Uttar Pradesh also showed Karp as the predominant genotype (15–17).

The common method used for genotyping in India is nested PCR amplification of partial gene followed by sequencing using protocol described by Furuya and Horinouchi (6, 30). In Furuya protocol, the amplified segment has 418-453 nucleotides which covers two variable domain (VD II and VD III) whereas the Horinouchi protocol covers three variable domains (VD I to III) and has ≈652 nucleotides in the whole 1600 nucleotides thus simplifying the whole process (6,18). The partial sequences of the TSA 56 gene will not provide full information about the entire ORF. For accurate classification and analysis, sequencing of entire ORF (including VD I to VD IV) of the 56 kDa is needed (9,19). As of date, there are 90,716 partial gene sequences available and only 609 complete gene sequences available in the GenBank (https://www.ncbi.nlm.nih.gov/nuccore?term=tsa56+complete+gene&cmd). The Indian genotype data is based on partial gene sequence of the TSA 56 gene and there is no complete gene 56 kDa data till date (Prakash JA Personal communication). We undertook a study to amplify the complete TSA 56 gene to definitely determine the circulating *Orientia tsutsugamushi* genotypes in and around Vellore. Further, we compared the complete TSA 56 gene data with the partial gene sequences obtained using the Furuya and Horinouchi protocol.

## Materials and methods

### Sample collection, processing and DNA extraction

Patients of either sex, above 1 year of age with fever more than 3 days less than 10 days with or without eschar/rash were recruited for this study after obtaining informed consent (EC approval no. 11942 dated 27^th^ March, 2019) from December 2020 to November 2021. Whole blood (8 ml) was collected in cell processing tubes (CPT) with sodium heparin (BD Vacutainer® CPT™ Franklin lakes, NJ, USA) from all study patients. The collected samples were centrifuged at 1600 RCF for 20 minutes at room temperature. The centrifugation resulted in the gel separating the mononuclear cells in the plasma from the erythrocytes and granulocytes. The separated plasma was aliquoted into two tubes; the monocyte layer was pipetted carefully and aliquoted into four tubes with the addition of equal volume of 40% glycerol. The plasma and PBMC aliquots were stored at -70ºC. One aliquot was used for DNA extraction using the Promega Wizard® Genomic DNA Purification kit (Promega Corporation, Madison, WI, USA) as per the manufacturer’s protocol. The extracted DNA was quantified using Nanodrop spectrophotometer (ThermoFisher Scientific, Waltham, MA, USA) and stored at -80ºC for pending molecular assays.

### Molecular assays performed

#### Real time PCR for 47 kDa gene

The extracted DNA was subjected to 47 kDa real-time PCR on an Applied Biosystems™ 7500 Real-Time PCR System (Thermo Fisher Scientific, MA, USA). This real-time PCR amplifies a 118bp section of 47 kDa gene (htrA) using the primers OtsuFP630 and OtsuRP747 and the probe OtsuPR665 as described previously (6). The 47kDa target gene amplification was done using TaqMan Fast Advanced Mastermix (Thermo Fisher Scientific, Waltham, MA, USA). Each reaction mixture (25 µl) contained 5 µl of template DNA, 10 pmol of each primer (10 pmol each), 12.5 µL of Master Mix and 5 pmol of the probe. The PCR conditions included an initial denaturation at 95ºC for 5 minutes followed by 40 cycles of 95°C for 30 seconds and 60°C for 1 minute. The Ct value ≤ 38 was considered as positive (13).

### Conventional PCR

Each PCR amplification method was performed in a 25 µl reaction volume using HotStar Taq Plus Master Mix kit (Qiagen, Hilden, Germany). Details of primers used in this study are given in the **Table 1**.

**Table 1:**
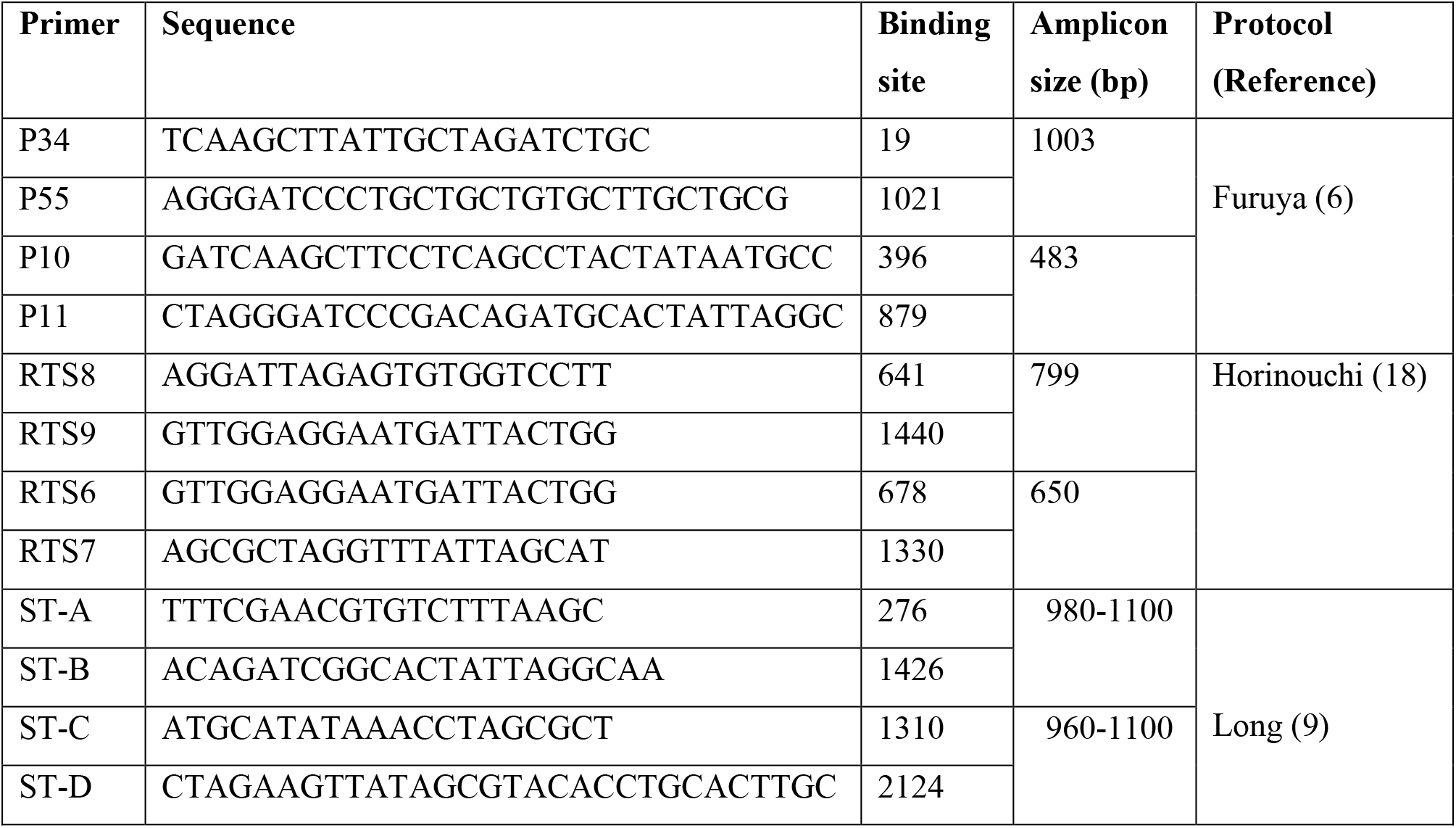
Details of primers used for 56kDa gene amplification.

#### Amplification of complete 56kDa gene as two fragment

The samples which had 47 kDa real-time PCR Ct values ≤ 30 were selected to amplify TSA 56 gene complete ORF (1605 bp). The TSA 56 gene was amplified using two pairs of primers as described by Long *et al*., in 2020 (9). The primer pair ST-A and ST-B amplified the first half of the TSA 56 gene. The reaction mixture contained 12.5 µl of Master Mix and 10 pmol of each primer with the PCR conditions as follows: 95ºC for 5minutes (for the activation of polymerase); 35 cycles of denaturation, annealing and extension at 94ºC for 1 minute, 55ºC for 1 minute 30 seconds, 72ºC for 2 minutes with a final extension step of 72ºC for 10 minutes. The primer pair ST-C and ST-D was used to amplify second half of the TSA 56 gene with the same PCR conditions as for the first half except the annealing time, which was shortened to 1 minute. The amplified fragments were analysed by agarose gel electrophoresis and visualized by gel documentation system (Gel Doc, Bio-Rad, Hercules, California, USA).

#### Sanger sequencing

The sample positive for both the fragment (first half and second half) were selected for sequencing. The amplified fragment were subjected to pre-clean up using Exosap-IT Applied Biosystems (ThermoFisher Scientific, Waltham, MA, USA) method with reaction conditions as follows: 37ºC for 15 minutes, 80ºC for 15 minutes and 15 ºC for 2 minutes. The pre-clean up product was used as template for sequencing PCR to amplify individual strands of the fragments BigDye Terminator v3.1 Cycle Sequencing Kit (Applied Biosystems, Foster City, CA, USA). The amplified segment was subjected to post cleanup using HighPrep™ DTR Clean-up System (MagBio Genomics Inc., Gaithersburg, MD, USA) and eluted with molecular grade water. The eluted product was loaded in Genetic Analyzer 3500 (ThermoFisher Scientific, Waltham, MA, USA)] and the raw reads was analysed and spliced using Bio-Edit software (20).

### Amplification of Complete TSA 56 gene as single fragment

Sample which showed poor quality reads were amplified using ST-A (forward primer of first fragment) and ST-D (reverse primer of second fragment). The reaction mixture contained 12.5 µl of Master Mix and 10 pmol of each primer with the PCR conditions as follows: 95ºC for 5minutes (for the activation of polymerase); 35 cycles of denaturation, annealing and extension at 94ºC for 1 minute, 55ºC for 1minute 30 seconds, 72ºC for 2 minutes with a final extension step of 72ºC for 10 minutes. The amplified fragments were analysed by agarose gel electrophoresis and visualized by gel documentation system (GelDoc, Bio-Rad, Hercules, USA). The 1800 bp positive amplicons was purified using Wizard® SV Gel and PCR Clean-Up System (Promega Corporation, Madison, WI, USA).

### Nested PCR for amplification of partial 56kDa gene

Samples positive for complete gene amplification were subjected to a nested PCR protocol as described by Horinouchi. This amplifies a 650 bp segment of the TSA 56 gene (includes VD-1 to VD-III) and used recently by Masakhwe *et al*., 2018 (18). The PCR product was analysed using agarose gel electrophoresis system and visualized by gel documentation (Gel Doc, Bio-Rad, Hercules, California, USA). The 650 bp positive fragment was purified by Wizard® SV Gel and PCR Clean-Up System (Promega Corporation, Madison, WI, USA).

Further, we performed a nested PCR to amplify a 483 bp fragment, encompassing VD-II and VD-III segment of the same gene, described by Furuya which were extensively evaluated by Kim *et al*., (6). As the PCR amplified product obtained using Furuya protocol showed multiple bands, we performed *in-silico* analysis for determining the genotype.

### Next Generation Sequencing

Totally 18 amplicons (11 partial and 7 complete) were sequenced on GridION X5 (Oxford Nanopore Technologies, Oxford, UK) using Spot ON flow cell R9.4 (FLO-MINI06) in a 48 hrs sequencing protocol. The resulting raw data showed good sequencing coverage and high read depth. High quality processed data was aligned against reference sequence (Karp M33004) and the consensus sequence was generated.

### In-silico analysis of partial gene

To obtain 483 bp fragment, the 9 complete and 4 near complete sequences were aligned with the inner primer P10 and P11 (of the furuya protocol) using CLUSTAL Omega sequence alignment tool (21). The inner primer covers the VD II and VD III region was selected and subjected to phylogenetic analysis. The two samples (CMCOT1 and CMCOT7) which showed low amplification by Horinouchi protocol were also subjected to *in-silico* generation of fragments by aligning the respective complete sequence with inner primer RTS 6 and RTS 7.

#### Phylogenetic tree

Both Sanger and NGS sequence was subjected to BLAST analysis. Alignment was performed using Clustal Omega (21) with 37 reference sequences retrieved from GenBank-NCBI and the phylogenetic tree was established separately for complete and partial gene using IQTREE software: A fast and effective stochastic algorithm for estimating maximum likelihood phylogenies (22).

### Analysis of Open Reading Frame (ORF) of complete TSA 56 gene

The ORF of the 13 sequences was predicted using NCBI Open Reading Frame Finder: RRID:SCR_016643 (23). The variable domains of the amino acid sequence were identified using Clustal Omega: RRID:SCR_001591 (21) alignment along with the respective reference sequence (Karp, Gilliam, Kato and TA763) (Supplementary Figure 1)

## Results

A total of 379 blood samples of scrub typhus suspected patient were collected from December 2020 to November 2021. Among these 162 samples were positive for 47 kDa real time PCR with Ct value ≤ 38 of which 73 samples had Ct value ≤ 30. Amongst 21 positive samples tested the Long protocol provided 9 complete and 4 near complete sequences (totally 13). Of these, 6 were Sanger derived sequences while other 7 were Nanopore derived.

Among the 6 Sanger sequenced amplicons, 2 had complete ORF (1605 bp) whereas in 4 the length varied from 1551 to 1596 bp. All the seven Nanopore sequenced samples showed complete ORF.

The Horinouchi protocol amplified 11 samples from the 13 tested generated ≈650 bp sequence. All the sequences have been submitted to the GenBank and the accession number received are as follows: 56kda complete/near complete sequence: OK545864, OK642783-86, OL631134, and OP037795-801; 56kDa partial (≈650bp) sequence: OP068199-209.

The BLAST result of complete gene sequences CMCOT-1 to CMCOT-13 showed 96.48 to 98.53% homology with *Orientia tsutsugamushi* strains AY787232, GQ332743, MW495499, GU120144, GU120140, GU446592, GU120150, GU446599, GU120146, GU446600, MW495814. The partial gene 650bp sequences showed 98.32 to 99.85℅ homology to KR073080, KR073078, KR073079, MT258805, KR073077, KR073076 strains. The phylogenetic tree was constructed separately for partial and complete gene using 37 standard reference sequences retrieved from GenBank. The ORF of the reference sequences ranged from 1422 bp to 1608 bp.

Phylogenetic analysis of the 13 complete/near complete 56kDa gene sequenced showed that 12 clustered into TA763B genotype and TA763 genogroup whereas one sample belongs to Karp A genotype and Karp genogroup (**Figure 2a**). For the Horinouchi protocol (650bp) we had 11 amplicons derived sequences and 2 *in-silico* sequences derived from the complete gene (Long protocol). Phylogenetic analysis revealed results similar to that obtained for the complete/near complete gene analysis i.e. 12 belong to TA763 and 1 belongs to Karp genotype (**Figure 2b**). For the 13 *in-silico* Furuya sequences, 10 belonged to TA763 genotype whereas the remaining 3 fall in the Karp genotype (**Figure 2c**). Supplementary Table 1 summarizes the phylogenetic analysis described above.

**Figure 2a:**
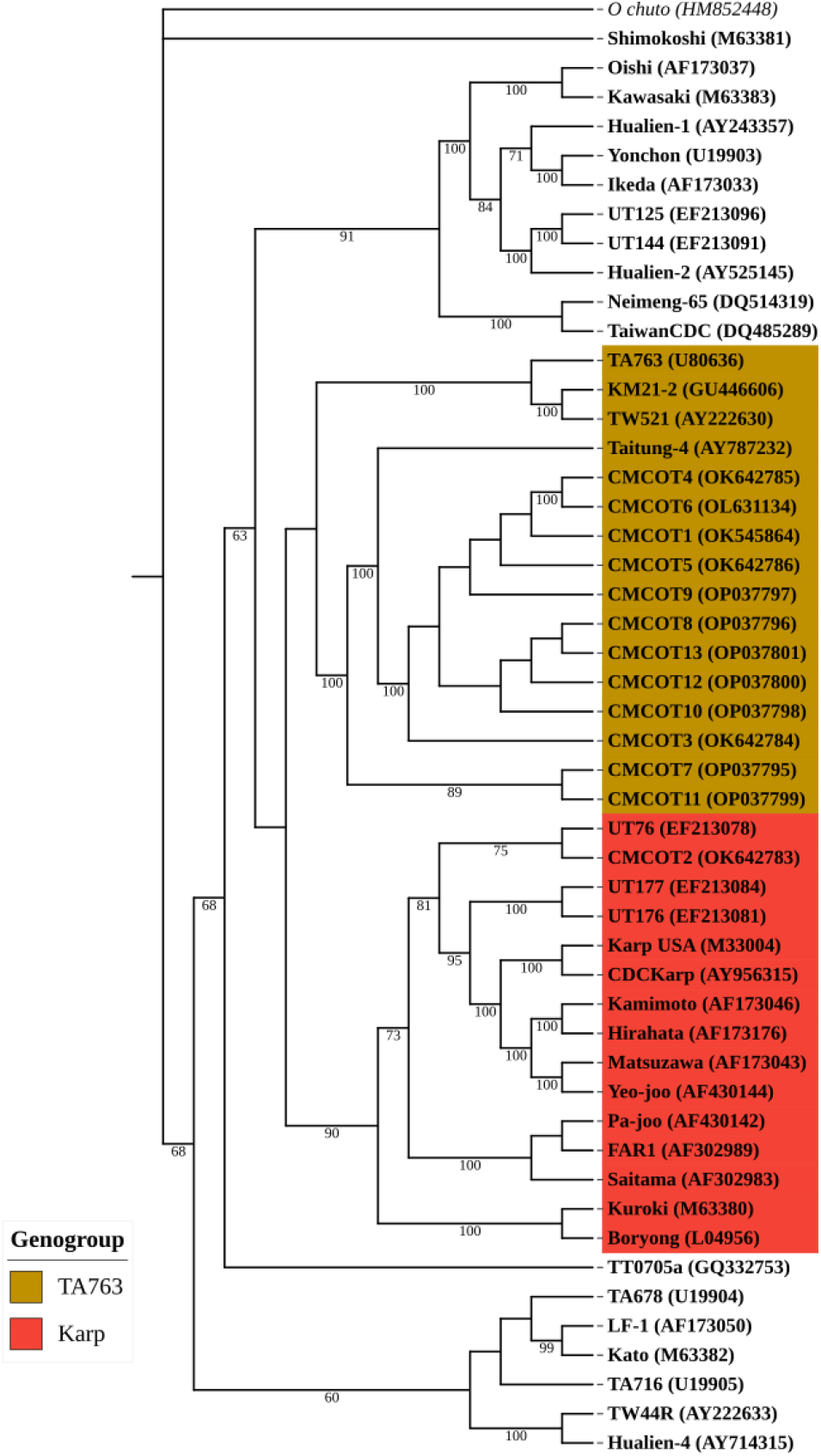
Phylogenetic tree of complete TSA 56 gene sequence: Long protocol.

**Figure 2b:**
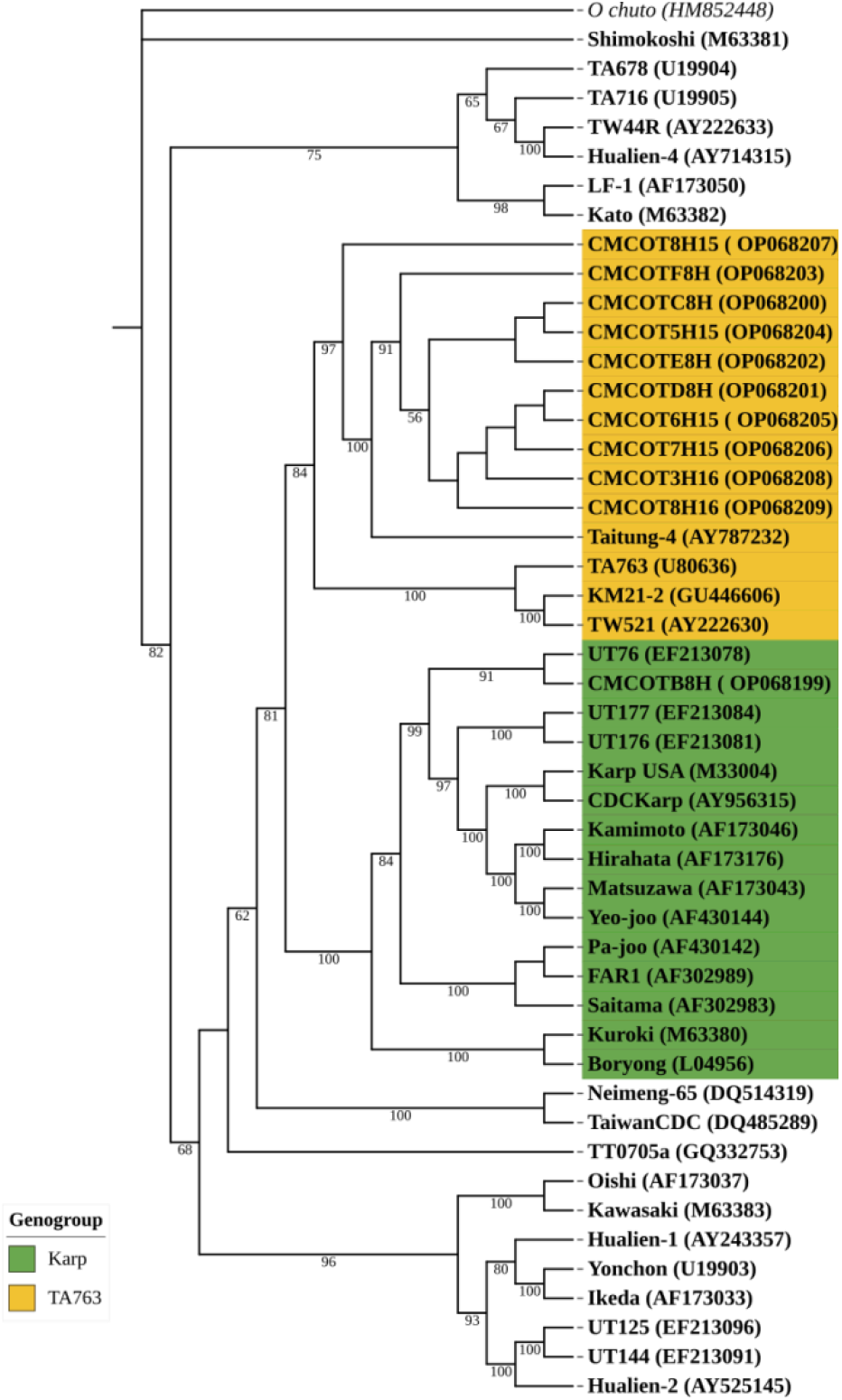
Phylogenetic tree of 650bp partial gene sequence: Horinouchi protocol.

**Figure 2c:**
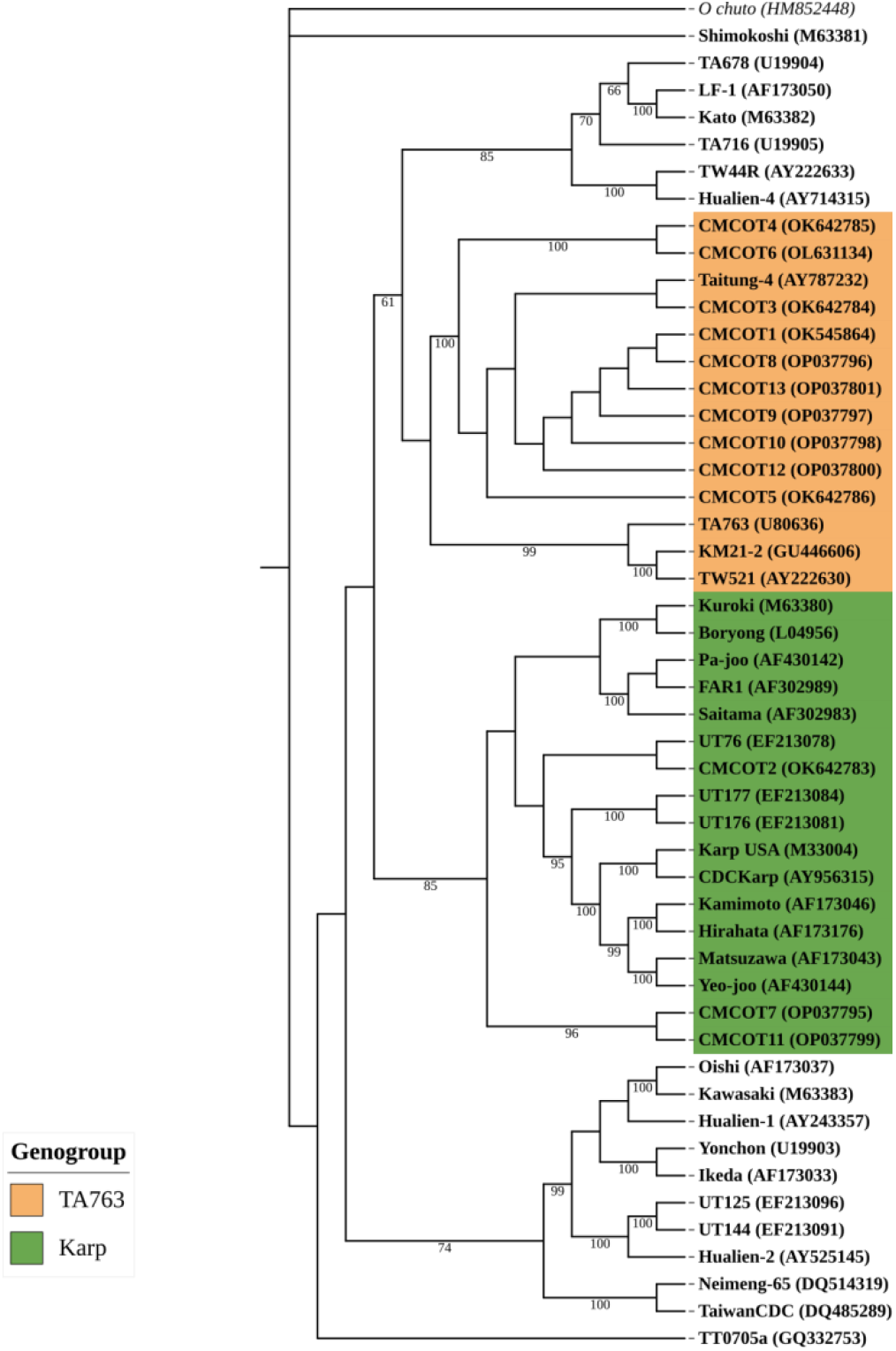
Phylogenetic tree of 480bp partial gene sequence: Furuya protocol.

## Discussion

Identification of *Orientia tsutsugamushi* genotypes circulating in endemic areas is important for designing region specific diagnostic kits and vaccines. This is because immunity is genotype specific (7, 8). The genotype variation is due to the variations in the four variable domains (VD I-IV) of the immuno dominant type specific TSA 56 gene. Of these, VD I-IV is the most variable and is the most important in determining the genotype (12). Knowledge of the circulating genotypes in a given area requires amplification of the complete gene followed by sequencing and phylogenetic analysis (6, 10, 30, 31). In India, such data is not available, we present the first comprehensive genotyping data based on amplification and sequencing of the four variable domains (VD I-IV) of the immuno dominant type specific 56 kDa gene.

In this study, we amplified the complete 56 kDa gene (≈1600bp) from 21 samples of which only 13 could be successfully sequenced. Point mutations and recombination are quite common in the variable regions which adversely affect the primer annealing and sensitivity of PCR (24). We compared three protocols for 56 kDa gene amplification. The Long protocol covers four variable domains (VD-I to VD-IV). The Horinouchi protocol covers three variable domains (VD-I to VD-III) and the Furuya protocol covers VD-II and VD-III regions (6,9,18). The complete gene was successfully amplified in 13 samples and fidelity of amplification was confirmed as more than 96% sequence homology was obtained with *Orientia tsutsugamushi* 56 kDa complete gene sequences by BLAST. Of the 6 samples sequenced using Sanger sequencing method, Only 2 were complete (1605 bp) whereas others are falling short of 10-50 nucleotides at 3’ end. This may be due to the binding region of ST-D primer was exactly at the end of the ORF (9). The first 20 to 40 bases are typically not well resolved in Sanger sequencing similarly the end region does not have well defined peaks. For optimal sequence data, it is important to design primer 60-100 bp away from the target region. Further, Sanger sequencing provides best peak resolution between 100 to 500 bases (25). Considering these limitations, the 7 sample showing poor quality reads in Sanger sequencing were again amplified as single fragment (1800 bp) using ST-A and ST-D primer and sequenced using Nanopore technology. The amplified 650 bp partial gene also sequenced using Nanopore technology along with the complete sequence. The 480 bp partial sequence (Furuya) showed multiple bands which may be due to non-specific binding of primer to the DNA (26). However, we obtained the 480 bp fragment for the 13 sample by *in-silico* method using the complete gene sequence.

Phylogenetic analysis of complete and partial gene revealed the genotypes as TA763 and Karp. The complete and 650 bp partial 56 kDa gene provides similar result whereas the 480 bp partial gene provides discordant result for two sequences (CMCOT7 and CMCOT11). Based on complete and 650 bp partial gene phylogeny CMCOT7 and CMCOT11 belongs to TA763 genotype but 480 bp partial gene phylogeny shows that the two sequences belong to Karp genotype. This discrepancy is due to the 480 bp partial gene covers only 2 variable domains (VDII and VDIII) whereas the 650bp sequence covers three variable domains (VDI to VDIII). The variable domain IV doesn’t play much role as the variations in the sequence are mostly conserved or semi-conserved (7). Therefore, the 650bp partial gene (Horinouchi protocol) is enough to determine the genotype of *Orientia tsutsugamushi* as it provides same result as complete gene analysis. Sequencing the Horinouchi protocol derived amplicons can be done by Sanger sequencing, which has good fidelity for amplification up to 800 bp (27,28).

Our limited data suggests that, TA763 is the predominant genotype circulating in Vellore. This genotype has been reported in South East Asia including Thailand and other countries like China, Australia and Taiwan (28,29). Only one sequence was found to be Karp. Globally, Karp genotype is reported to account for about 39.5% and found throughout the endemic region (31). Our present study provides first report on *Orientia tsutsugamushi* genotypes using complete 56 kDa gene sequence. Multi-centric studies which include other genotypes are needed for validating this preliminary finding.

## Conclusion

TA763 is the commonest genotype based on phylogenetic analysis of the complete TSA 56 gene. Sequence analysis of the 650 bp gene fragment encompassing the variable regions VDI-III is a promising method for detecting the genotype. Moreover, the shorter 650 bp amplicons generated can be easily sequenced by the Sanger protocol.

## Acknowledgements

None

## Author contributions

Conceptualization and methodology: K.G., A.K.P.P., J.A.J.P.; Investigations and Data acquisition: J.K., P.G., L.S.N.; Analysis and data interpretation: J.K., P.G., L.S.N., J.A.J.P., Drafting the manuscript: J.K., L.S.N.; Critical revision of the manuscript: K.G., A.K.P.P., J.A.J.P.; Supervision and funding acquisition: J.A.J.P.

