## Supplementary Figure 1 for "Genotyping of *Orientia tsutsugamushi* circulating in and around Vellore (South India) using TSA 56 gene"

**Figure 1. Variable domains of 56 kDa gene sequences: Kato, Karp, Gilliam, TA763 and CMCOT 1 to 13**

```
KATO         MKKIMLIASAMSALSLPFSASAIELGDEGGLECGPYAKVGVVGGMITGVESTRLDPADAG	60
Gilliam      MKKIMLIASAMSALSLPFSASAIELGEEGGLECGPYGKVGIVGGMITGAESTRLDSTDSE	60
TA763        MKKIMLIASAMSALSLPFSASAIELGDEGGLECGPYAKVGVIGGMITGVESARLDPTDSE	60
CMCOT4       MKKIMLIASAMSALSLPFSASAIEMGDEGGLECGPYAKVGIVGGMITGAESTRLDSSDAE	60
CMCOT6       MKKIMLIASAMSALSLPFSASAIEMGDEGGLECGPYAKVGIVGGMITGAESTRLDSSDAE	60
CMCOT3       MKKIMLIASAMSALSLPFSASAIELGDEGGLECGPYAKVGIVGGMITGAESTRLDSSDAE	60
CMCOT1       MKKIMLIASAMSALSLPFSASAIELGDEGGLECGPYAKVGIVGGMITGAESTRLHSSDAE	60
CMCOT5       MKKIMLIASAMSALSLPFSASAIELGDEGGLECGPYAKVGIVGGMITGAESTRLDSSDAE	60
CMCOT8       MKKIMLIASAMSALSLPFSASAIELGDEGGLECGPYAKVGIVGGMITGAESTRLDSSDAE	60
CMCOT9       MKKIMLIASAMSALSLPFSASAIELGDEGGLECGPYAKVGIVGGMITGAESTRLDSSDAE	60
CMCOT10      MKKIMLIASAMSALSLPFSASAIELGDEGGLECGPYAKVGIVGGMITGAESTRLDSSDAE	60
CMCOT12      MKKIMLIASAMSALSLPFSASAIELGDEGGLECGPYAKVGIVGGMITGAESTRLDSSDAE	60
CMCOT13      MKKIMLIASAMSALSLPFSASAIELGDEGGLECGPYAKVGIVGGMITGAESTRLDSSDAE	60
CMCOT7       MKKIMLIASAMSALSLPFSASAIELGDEGGLECGPYAKVGVVGGMITGVESARLDPADAD	60
CMCOT11      MKKIMLIASAMSALSLPFSASAIELGDEGGLECGPYAKVGVVGGMITGVESARLDPADAD	60
KARP         MKKIMLIASAMSALSLPFSASAIELGEE-GLECGPYAKVGVVGGMITGVESARLDPADAE	59
CMCOT2       MKKIMLIASAMSALSLPFSASAIELGDEGGLECGPYAKVGVVGGMITGVESARLDPADAD	60
             ************************:*:* *******.***::******.**:**. :*:

KATO         GKKQLPLTTSMPFGGTLAAGMTIAPGFRAELGVMYLANVKAEVESGKTGSDADIRS----	116
Gilliam      GKKHLSLTTGLPFGGTLAAGMTIAPGFRAELGVMYLRNISAEVEVGKGKVDSKGEIKADS	120
TA763        GKKHLSLTTGMPFGGTLAAGMTITPSIRAELGVMYLRNISAEVELGKVKADSGSKTKADS	120
CMCOT4       GKKRLSLTTSMPFGGTLAAGMTIAQGFRAELGVMYLTNITAQVEEGKGKIGSNVAKGSDA	120
CMCOT6       GKKRLSLTTSMPFGGTLAAGMTIAQGFRAELGVMYLTNITAQVEEGKGKIGSNVAKGSDA	120
CMCOT3       GKKRLSLTTSMPFGGTLAAGMTIAQGFRAELGVMYLTNITAQVEEGKGKIGSNVAKGSDA	120
CMCOT1       GKKRLSLTTSMPFGGTLAAGMTIAQGFRAELGVMYLTNITAQVEEGKGKIGSNVAKGSDA	120
CMCOT5       GKKRLSLTTSMPFGGTLAAGMTIAQGFRAELGVMYLTNITAQVEEGKGKIGSNVAKGSDA	120
CMCOT8       GKKRLSLTTSMPFGGTLAAGMTIAQGFRAELGVMYLTNITAQVEEGKGKIGSNVAKGSDA	120
CMCOT9       GKKRLSLTTSMPFGGTLAAGMTIAQGFRAELGVMYLTNITAQVEEGKGKIGSNVAKGSDA	120
CMCOT10      GKKRLSLTTSMPFGGTLAAGMTIAQGFRAELGVMYLTNITAQVEEGKGKIGSNVAKGSDA	120
CMCOT12      GKKRLSLTTSMPFGGTLAAGMTIAQGFRAELGVMYLTNITAQVEEGKGKIGSNVAKGSDA	120
CMCOT13      GKKRLSLTTSMPFGGTLAAGMTIAQGFRAELGVMYLTNITAQVEEGKGKIGSNVAKGSDA	120
CMCOT7       GKKQLPLTTSMPFGGTLAAGMTIAPGFRAELGVMYLTNITAQVEEGKVKADSNVDKGSDS	120
CMCOT11      GKKQLPLTTSMPFGGTLAAGMTIAPGFRAELGVMYLTNITAQVEEGKVKADSEVDKGSDS	120
KARP         GKKHLSLTNGLPFGGTLAAGMTIAPGFRAEIGVMYLTNITAQVEEGKVKADSVGETKADS	119
CMCOT2       GKKQLPLTTSMPFGGTLAAGMTIAPGFRAELGVMYLTNITAQVEEGKVKADSGGKTKADS	120
             ***:* **..:************: .:***:***** *:.*:** **   .:

KATO         ---------GADSPMPQRYKLTPPQPTIMPISIADRDLGVDIPNVPQGGANHLGDNLGAN	167
Gilliam      GG-------GTDTPIRKRFKLTPPQPTIMPISIADRDVGVDTDILAQAAA--GQ--PRLT	169
TA763        GG-------ETDAPIRKRFKLTPPQPTIMPISIADRDFGVDVTNIPQAQVQPPQ--QAND	171
CMCOT4       ANDNTGTDANQAKQLKRPPKLTPPQPVIMPISTADRDMGVDVINEPQAQLAQAQ--Q-ND	177
CMCOT6       ANDNTGTDANQAKQLKRPPKLTPPQPVIMPISTADRDMGVDVINEPQAQLAQAQ--Q-ND	177
CMCOT3       ANDNTGTDANQAKQLKRPPKLTPPQPVIMPISTADRDMGVDVINEPQAQVAQAQ--Q-ND	177
CMCOT1       ANDNTGTDANQAKQLKRPPKLTPPQPVIMPISTADRDMGVDVINEPQAQVAQAQ--Q-ND	177
CMCOT5       ANDNTGTDANQAKQLKRPPKLTPPQPVIMPISTADRDMGVDVINEPQAQVAQAQ--Q-ND	177
CMCOT8       ANDNTGTDANQAKQLKRPPKLTPPQPVIMPISTADRDMGVDVINEPQAQVAQAQ--Q-ND	177
CMCOT9       ANDNTGTDANQAKQLKRPPKLTPPQPVIMPISTADRDMGVDVINEPQAQVAQAQ--Q-ND	177
CMCOT10      ANDNTGTDANQAKQLKRPPKLTPPQPVIMPISTADRDMGVDVINEPQAQVAQAQ--Q-ND	177
CMCOT12      ANDNTGTDANQAKQLKRPPKLTPPQPVIMPISTADRDMGVDVINEPQAQVAQAQ--Q-ND	177
CMCOT13      ANDNTGTDANQAKQLKRPPKLTPPQPVIMPISTADRDMGVDVINEPQAQVAQAQ--Q-ND	177
CMCOT7       ANDNTGTDANQAKQRKRPPKLTPPQPTIMPISIADRDLGVDICNIPQAQVAQAQ--Q-ND	177
CMCOT11      GKDKTGTDAXXAILRKRLFKLTPPQPTIMPISIADRDLGIDICNIPQAQLAQAP--H-ND	177
KARP         ---VGGKD----APIRKRFKLTPPQPTIMPISIADRDFGIDIPNIPQQQAQAAQ--PQLN	170
CMCOT2       ---GGGTD----APIRKRFKLTPPQPTIMPISIADRDFGIDICNIPHAQAQAAN--PALN	171
                             :  *******.***** ****.*:*     :

KATO         DIRRADDRITWLKNYAGVDYMVPDPNNPQA-RIVNPVLLNIPQGPPNANPR-----QAMQ	221
Gilliam      VEQRAADRIAWLKNYAGIDYMVPDPQNPNA-RVINPVLLNITQGPPNVQPRP------RQ	222
TA763        PLVRGVRRIAWLKEYAGIDYMVKDPNNP-GRMMVNPVLLNIPQGPPAQNPR-----AAMQ	225
CMCOT4       PLVRGLRRIAWLKQYAGIDYMVKDPNNP-GQMMVNPVLLNIPQGPPANNPR-----APMQ	231
CMCOT6       PLVRGLRRIAWLKQYAGIDYMVKDPNNP-GQMMVNPVLLNIPQGPPANNPR-----APMQ	231
CMCOT3       PLVRGLRRIAWLKQYAGIDYMVKDPNNP-GQMMVNPVLLNIPQGPPANNPR-----APMQ	231
CMCOT1       PLVRGLRRIAWLKQYAGIDYMVKDPNNP-GQMMVNPVLLNIPQGPPANNPR-----APMQ	231
CMCOT5       PLVRGLRRIAWLKQYAGIDYMVKDPNNP-GQMMVNPVLLNIPQGPPANNPR-----APMQ	231
CMCOT8       PLVRGLRRIAWLKQYAGIDYMVKDPNNP-GQMMVNPVLLNIPQGPPANNPR-----APMQ	231
CMCOT9       PLVRGLRRIAWLKQYAGIDYMVKDPNNP-GQMMVNPVLLNIPQGPPANNPR-----APMQ	231
CMCOT10      PLVRGLRRIAWLKQYAGIDYMVKDPNNP-GQMMVNPVLLNIPQGPPANNPR-----APMQ	231
CMCOT12      PLVRGLRRIAWLKQYAGIDYMVKDPNNP-GQMMVNPVLLNIPQGPPANNPR-----APMQ	231
CMCOT13      PLVRGLRRIAWLKQYAGIDYMVKDPNNP-GQMMVNPVLLNIPQGPPANNPR-----APMQ	231
CMCOT7       PLVRASPRIAWLKNCAGIDYRVKDPNNP-GPMVINPILLNIPQGPPAGNPR-----APMQ	231
CMCOT11      PLVRAAARIAWLKNCAGIDYRVKDPNNP-GPMVINPILLNIPQGPPAGNPR-----APMQ	231
KARP         DEQRAAARIAWLKNCAGIDYRVKNPNDPNGPMVINPILLNIPQGNPNPVGNPPQRANPPA	230
CMCOT2       DEQRAAARIAWLKNCAGIDYRVKDPNNPNGPMVINPILLNIPQGNPNPAGNPPQRAQQPA	231
                *.  **:***: **:** * :*::* .  ::**:**** ** *    .

KATO         PCSILNHDHWRHLVVGITAMSNANKPSVSPIKVLSEKIVQIYRDVKPFARVAGIEVPSDP	281
Gilliam      NLDILDHGQWRHLVVGVTALSHANKPSVTPVKVLSDKITKIYSDIKPFADIAGIDVPDTG	282
TA763        PCNILDHDHWKHFVVGVTALSNANKPSASPVKILSEKITQIYSDIRPFADIAGIDVPDAG	285
CMCOT4       RCDILNHDHWRHLLVGLTASSHPPKPSASPLKALTDKITQIYSDIKPFADIAGIDVPDTG	291
CMCOT6       RCDILNHDHWRHLLVGLTASSHPPKPSASPLKALTDKITQIYSDIKPFADIAGIDVPDTG	291
CMCOT3       RCDILNHDHWRHLVVGIAALSNANKPSASPVKVLSDKITQIYSDIKPFADIAGIDVPDTG	291
CMCOT1       RCDILNHDHWRHLVVGIAALSNANKPSASPVKVLSDKITQIYSDIKPFADIAGIDVPDTG	291
CMCOT5       RCDILNHDHWRHLVVGIAALSNANKPSASPVKVLSDKITQIYSDIKPFADIAGIDVPDTG	291
CMCOT8       RCDILNHDHWRHLVVGIAALSNANKPSASPVKVLSDKITQIYSDIKPFADIAGIDVPDTG	291
CMCOT9       RCDILNHDHWRHLVVGIAALSNANKPSASPVKVLSDKITQIYSDIKPFADIAGIDVPDTG	291
CMCOT10      RCDILNHDHWRHLVVGIAALSNANKPSASPVKVLSDKITQIYSDIKPFADIAGIDVPDTG	291
CMCOT12      RCDILNHDHWRHLVVGIAALSNANKPSASPVKVLSDKITQIYSDIKPFADIAGIDVPDTG	291
CMCOT13      RCDILNHDHWRHLVVGIAALSNANKPSASPVKVLSDKITQIYSDIKPFADIAGIDVPDTG	291
CMCOT7       XFAIHNHDHWRHLVVGLAALSNANKPSASPVKVLSDKITQIYSDIKPFADIAGIDVPDTS	291
CMCOT11      IFAIHNHDHWRHLVVGLAALSNANKPSASPVKVLSDKITQIYSDIKPFADIAGIDVPDTS	291
KARP         GFAIHNHEQWRHLVVGLAALSNANKPSASPVKVLSDKITQIYSDIKPFADIAGIDVPDTS	290
CMCOT2       NFAIHNHDHCRHLVVGLAALSNANKPSASPVKVLSDKITQIYSDIKPFADIAGIDVPDTG	291
                * :* : :*::**::* *:  ***.:*:* *::**.:** *::*** :***:**.

KATO         LPNSASVEQIQNKMQELNDILDEIRDSFDGCIGGNAFANQIQLNFRIPQAQQQGQGQ-QQ	340
Gilliam      LPNSASVEQIQSKMQELNDVLEDLRDSFDGYMG-NAFANQIQLNFVMPQQAQQQQGQGQQ	341
TA763        LPNSATVEQIQNKMQELNDVLEELRESFDGYLGGNAFANQIQLNFVMPQQA-QQQGQGQQ	344
CMCOT4       LPNSASVEQIQRKMQELNDVLEGLRDAFDGYIN-NAFVDQIQLNFVMPPQAQQQQGQGQQ	350
CMCOT6       LPNSASVEQIQRKMQELNDVLEGLRDAFDGYIN-NAFVDQIQLNFVMPPQAQQQQGQGQQ	350
CMCOT3       LHNSASVEQIQRKMQELNDVLEGLRDAFDGYIN-NAFVDQIQLNFVMPPQAQQQQGQGQQ	350
CMCOT1       LPNSASVEQIQRKMQELNDVLEGLRDAFDGYIN-NAFVDQIQLNFVMPPQAQQQQGQGQQ	350
CMCOT5       LPNSASVEQIQRKMQELNDVLEGLRDAFDGYIN-NAFVDQIQLNFVMPPQAQQQQGQGQQ	350
CMCOT8       LPNSASVEQIQRKMQELNDVLEGLRDAFDGYIN-NAFVDQIQLNFVMPPQAQQQQGQGQQ	350
CMCOT9       LPNSASVEQIQRKMQELNDVLEGLRDAFDGYIN-NAFVDQIQLNFVMPPQAQQQQGQGQQ	350
CMCOT10      LPNSASVEQIQRKMQELNDVLEGLRDAFDGYIN-NAFVDQIQLNFVMPPQAQQQQGQGQQ	350
CMCOT12      LPNSASVEQIQRKMQELNDVLEGLRDAFDGYIN-NAFVDQIQLNFVMPPQAQQQQGQGQQ	350
CMCOT13      LPNSASVEQIQRKMQELNDVLEGLRDAFDGYIN-NAFVDQIQLNFVMPPQAQQQQGQGQQ	350
CMCOT7       LPNSASVEQIQNKMQELNDVLEELRESFDGYLG-NAFADQIQLNFVMPPQAQQQQGQGQQ	350
CMCOT11      LPNSASVEQIQNKMQELNDVLEELRESFDGYLG-NAFADQIQLNFVMPPQAQQQQGQGQQ	350
KARP         LPNSASVEQIQNKMQELNDLLEELRESFDGYLGGNAFANQIQLNFVMPQQA-QQQGQGQQ	349
CMCOT2       LPNSASVEQIQRKMQELNDVLEGLRDAFDGYIN-NAFVDQIQLNFVMPPQAQQQQGQGQQ	350
             * ***:***** *******:*: :*::*** :. ***.:****** :*    * *** **

KATO         QQAQATAQEAAAAAAVRVLNNNDQIIKLYKDLVKLKRHAGIKKAMEELAAQDG---GCNG	397
Gilliam      QQAQATAQEAVAAAAVRLLNGNDQIAQLYKDLVKLQRHAGVKKAMEKLAAQQEEDAKNQC	401
TA763        QQAQATAQEAVAAAAVRLLNGNDQIAQLYRDLVKLQRHAGIKKAMEKLAAQQEEDAKNQG	404
CMCOT4       QQAQATAQEAVAAAAVRLLNGNDQIVQLYKDLVKLQRHAGIKKAMEKLAAQQEEDAKNQG	410
CMCOT6       QQAQATAQEAVAAAAVRLLNGNDQIVQLYKDLVKLQRHAGIKKAMEKLAAQQEEDAKNQG	410
CMCOT3       QQAQATAQEAVAAAAVRLLNGNDQIVQLYKDLVKLQRHAGIKKAMEKLAAQQEEDAKNQG	410
CMCOT1       QQAQATAQEAVAAAAVRLLNGNDQIVQLYKDLVKLQRHAGIKKAMEKLAAQQEEDAKNQG	410
CMCOT5       QQAQATAQEAVAAAAVRLLNGNDQIVQLYKDLVKLQRHAGIKKAMEKLAAQQEEDAKNQG	410
CMCOT8       QQAQATAQEAVAAAAVRLLNGNDQIVQLYKDLVKLQRHAGIKKAMEKLAAQQEEDAKNQG	410
CMCOT9       QQAQATAQEAVAAAAVRLLNGNDQIVQLYKDLVKLQRHAGIKKAMEKLAAQQEEDAKNQG	410
CMCOT10      QQAQATAQEAVAAAAVRLLNGNDQIVQLYKDLVKLQRHAGIKKAMEKLAAQQEEDAKNQG	410
CMCOT12      QQAQATAQEAVAAAAVRLLNGNDQIVQLYKDLVKLQRHAGIKKAMEKLAAQQEEDAKNQG	410
CMCOT13      QQAQATAQEAVAAAAVRLLNGNDQIVQLYKDLVKLQRHAGIKKAMEKLAAQQEEDAKNQG	410
CMCOT7       QQGQATAQEAVAAAAVRLLNGNDQIAQLYKDLVKLQRHAGIKKAMEKLAAQQEEDAKNQG	410
CMCOT11      QQGQATAQEAVAAAAVRLLNGNDQIAQLYKDLVKLQRHAGIKKAMEKLAAQQEEDAKNQG	410
KARP         QQAQATAQEAVAAAAVRLLNGNDQIAQLYKDLVKLQRHAGIKKAMEKLAAQQEEDAKNQG	409
CMCOT2       QQAQATAQEAVAAAAVRLLNGNDQIVQLYKDLVKLQRHAGIKKAMEKLAAQQEEDAKNQG	410
             **.*******.******:**.**** :**:*****:****:*****:****:      :

KATO         GGDNKKKRGASEDSDAGGASKGGKGKETKETEFDLSMIVGQVKLYADLFTTESFSIYAGL	457
Gilliam      EGDCKQQQGASEKSK--------EG-KGKETEFDLSMIVGQVKLYADLFTTESFSIYAGV	452
TA763        EGDCKQQQGTSEKSK--------EGS-KKEPEFDLSMIVGQVKLYADVMITESVSIYAGV	455
CMCOT4       EGDCKQQQGASEKSK--------EGGKGKEAEFDLSMIVGQVKLYADVMITESFSVYAGV	462
CMCOT6       EGDCKQQQGASEKSK--------EGGKGKEAEFDLSMIVGQVKLYADVMITESFSVYAGV	462
CMCOT3       EGDCKQQQRASEKSK--------EGGKGKDAEFDLSMIVGQVKLYADVMITESFSVYAGV	462
CMCOT1       EGDCKQQQGASEKSK--------EGGKGKEAEFDLSMIVGQVKLYADVMITESFSVYAGV	462
CMCOT5       EGDCKQQQGASEKSK--------EGGKGKEAEFDLSMIVGQVKLYADVMITESFSVYAGV	462
CMCOT8       EGDCKQQQGASEKSK--------EGGKGKEAEFDLSMIVGQVKLYADVMITESFSVYAGV	462
CMCOT9       EGDCKQQQGASEKSK--------EGGKGKEAEFDLSMIVGQVKLYADVMITESFSVYAGV	462
CMCOT10      EGDCKQQQGASEKSK--------EGGKGKEAEFDLSMIVGQVKLYADVMITESFSVYAGV	462
CMCOT12      EGDCKQQQGASEKSK--------EGGKGKEAEFDLSMIVGQVKLYADVMITESFSVYAGV	462
CMCOT13      EGDCKQQQGASEKSK--------EGGKGKEAEFDLSMIVGQVKLYADVMITESFSVYAGV	462
CMCOT7       EGDCKQQQGTSEKSK--------EGRKGKEAEFDLSMIVGQVKLYADLMTTESFSIYAGV	462
CMCOT11      EGDCKQQQGTSEKSK--------EGSKGKEAEFDLSMIVGQVKLYADLMTTESFSIYAGV	462
KARP         EGDCKQQQGTSEKSK--------K-GKDKEAEFDLSMIVGQVKLYADVMITESVSIYAGV	460
CMCOT2       EGDCKQQQGASEKSK--------EGGKGKEAEFDLSMIVGQVKLYADVMITESFSVYAGV	462
              ** *::: :**.*.        :    *: ****************:: ***.*:***:

KATO         GAGLAYTSGKIDGVDIKANTGMVASGALGVAINAAEGVYVDIEGSYMHSFSKIEEKYSIN	517
Gilliam      GAGLAHTYGKIDDKDIKGHTGMVASGALGVAINAAEGVYVDLEGSYMHSFSKIEEKYSIN	512
TA763        GAGLAYTSGKIDDKDTK-HTGMVVSGALGVAINAAEGVYVDIEGSYMYSFSKIEEKYSIN	514
CMCOT4       GAGLAYTYGKIDNKDIKGHTGMVASGALGVAINAAEGVYVDIEGGYMYSFSKIEEKYSIN	522
CMCOT6       GAGLAYTYGKIDNKDIKGHTGMVASGALGVAINAAEGVYVDIEGGYMYSFSKIEEKYSIN	522
CMCOT3       GAGLAYTYGKIHNNDIKGHTGMVASGALGVAINAAEGVYVDIQGGYMYSFSKIEE-----	517
CMCOT1       GAGLAYTYGKIDNKDIKGHTGMVASGALGVAINAAEGVYVDIEGGYMYSFSKIEEKYSIN	522
CMCOT5       GAGLAYTYGKIDNKDIKGHTGMVASGALGVAINAAEGVYVDIEGGYMYSFSKIEEKYSIN	522
CMCOT8       GAGLAYTYGKIDNKDIKGHTGMVASGALGVAINAAEGVYVDIEGGYMYSFSKIEEKYSIN	522
CMCOT9       GAGLAYTYGKIDNKDIKGHTGMVASGALGVAINAAEGVYVDIEGGYMYSFSKIEEKYSIN	522
CMCOT10      GAGLAYTYGKIDNKDIKGHTGMVASGALGVAINAAEGVYVDIEGGYMYSFSKIEEKYSIN	522
CMCOT12      GAGLAYTYGKIDNKDIKGHTGMVASGALGVAINAAEGVYVDIEGGYMYSFSKIEEKYSIN	522
CMCOT13      GAGLAYTYGKIDNKDIKGHTGMVASGALGVAINAAEGVYVDIEGGYMYSFSKIEEKYSIN	522
CMCOT7       GAGVAYTYGKIDNKDIKGHTGMVASGALGVAINAAEGVYVDIEGGYMHSFSKIEEKYSVN	522
CMCOT11      GAGVAYTYGKIDNKDIKGHTGMVASGALGVAINAAEGVYVDIEGGYMHSFSKIEEKYSVN	522
KARP         GAGLAYTSGKIDNKDIKGHTGMVASGALGVAINAAEGVYVDIEGSYMYSFSKIEEKYSIN	520
CMCOT2       GAGLAYTYGKIDNKDIKGHTGMVASRALGVAINAAEGVYVDIESGYMYSFSKIEEKYPIN	522
             ***:*:* ***.. * * :****.* ***************::..**:*******

KATO         PLMASFGVRYNF	529
Gilliam      PLMASVGVRYNF	524
TA763        PLMASVGVRYNF	526
CMCOT4       PLMASAGVRYNF	534
CMCOT6       PLMASAGVRYNF	534
CMCOT3       ------------	517
CMCOT1       PLMA--------	526
CMCOT5       PLMASAGVRY--	532
CMCOT8       PLMASAGVRYNF	534
CMCOT9       PLMASAGVRYNF	534
CMCOT10      PLMASAGVRYNF	534
CMCOT12      PLMASAGVRYNF	534
CMCOT13      PLMASAGVRYNF	534
CMCOT7       ALMASAGVRYNF	534
CMCOT11      ALMASAGVRYNF	534
KARP         PLMASVGVRYNF	532
CMCOT2       PLMASARV----	530
```
