## Supplementary Table 1 for "Genotyping of *Orientia tsutsugamushi* circulating in and around Vellore (South India) using TSA 56 gene"

| **Sample ID (Complete gene)** | Sequence size (in bp) | ORF (in bp) | Genotype (Complete gene) | **650bp Partial gene ID** | Sequence size (in bp) | Genotype (650bp Partial gene) | **480bp Partial gene ID (*in-silico*)** | Sequence size (in bp) | Genotype (480bp Partial gene) |
| --- | --- | --- | --- | --- | --- | --- | --- | --- | --- |
| CMCOT1 (OK545864) | 1832 | 1578 | TA763 | CMCOT1 H *(Insilico)* | 654 | TA763 | CMCOT1 F | 486 | TA763 |
| CMCOT2 (OK642783) | 1830 | 1590 | KARP | CMCOTB8H (OP068199) | 654 | KARP | CMCOT2 F | 507 | KARP |
| CMCOT3 (OK642784) | 1551 | 1551 | TA763 | CMCOTC8H (OP068200) | 654 | TA763 | CMCOT3 F | 486 | TA763 |
| CMCOT4 (OK642785) | 1773 | 1605 | TA763 | CMCOTD8H (OP068201) | 654 | TA763 | CMCOT4 F | 486 | TA763 |
| CMCOT5 (OK642786) | 1795 | 1596 | TA763 | CMCOTE8H (OP068202) | 654 | TA763 | CMCOT5 F | 486 | TA763 |
| CMCOT6 (OL631134) | 1773 | 1605 | TA763 | CMCOTF8H (OP068203) | 654 | TA763 | CMCOT6 F | 486 | TA763 |
| CMCOT7 (OP037795) | 1773 | 1602 | TA763 | CMCOT7 H *(Insilico)* | 654 | TA763 | CMCOT7 F | 486 | KARP |
| CMCOT8 (OP037796) | 1773 | 1602 | TA763 | CMCOT5H15 (OP068204) | 654 | TA763 | CMCOT8 F | 486 | TA763 |
| CMCOT9 (OP037797) | 1773 | 1602 | TA763 | CMCOT6H15 (OP068205) | 654 | TA763 | CMCOT9 F | 486 | TA763 |
| CMCOT10 (OP037798) | 1773 | 1602 | TA763 | CMCOT7H15 (OP068206) | 654 | TA763 | CMCOT10 F | 486 | TA763 |
| CMCOT11 (OP037799) | 1773 | 1602 | TA763 | CMCOT8H15 (OP068207) | 654 | TA763 | CMCOT11 F | 486 | KARP |
| CMCOT12 (OP037800) | 1773 | 1602 | TA763 | CMCOT3H16 (OP068208) | 654 | TA763 | CMCOT12 F | 486 | TA763 |
| CMCOT13 (OP037801) | 1773 | 1602 | TA763 | CMCOT8H16 (OP068209) | 654 | TA763 | CMCOT13 F | 486 | TA763 |

**Table 1: Genogroup assigned by phylogenetic analysis of complete and partial 56 kDa gene sequences**
